# AAV11 mediates efficient retrograde trans-neuronal transduction of astrocytes

**DOI:** 10.1101/2025.10.10.681765

**Authors:** Nengsong Luo, Yunling Gao, Zengpeng Han, Fuqiang Xu, Kunzhang Lin

**Affiliations:** Shenzhen Key Laboratory of Viral Vectors for Biomedicine, Shenzhen-Hong Kong Institute of Brain Science, Shenzhen Institutes of Advanced Technology, Chinese Academy of Sciences, 518055 Shenzhen, P. R. China; Wuhan National Laboratory for Optoelectronics, Huazhong University of Science and Technology, 430074 Wuhan, P. R. China; Key Laboratory of Magnetic Resonance in Biological Systems, State Key Laboratory of Magnetic Resonance and Atomic and Molecular Physics, National Center for Magnetic Resonance in Wuhan, Innovation Academy for Precision Measurement Science and Technology, Chinese Academy of Sciences, 430071 Wuhan, P. R. China; Key Laboratory of Quality Control Technology for Virus-Based Therapeutics, Guangdong Provincial Medical Products Administration, NMPA Key Laboratory for Research and Evaluation of Viral Vector Technology in Cell and Gene Therapy Medicinal Products, the Brain Cognition and Brain Disease Institute, Shenzhen Institutes of Advanced Technology, Chinese Academy of Sciences, 518055 Shenzhen, P. R. China; Institute of Neuroscience and Brain Diseases, Xiangyang Central Hospital, Affiliated Hospital of Hubei University of Arts and Science, 441106 Xiangyang, P. R. China; Department of Pediatrics, Boston Children’s Hospital, Harvard Medical School, 02115 Boston, MA, USA; Center for Excellence in Brain Science and Intelligence Technology, Chinese Academy of Sciences, 200031 Shanghai, P. R. China

**Author notes:** Correspondence (K.L.); (F.X.).

**Keywords:** astrocytes, AAV11, retrograde trans-neuronal tools, astrocyte-neuron networks

## Abstract

Understanding the mechanisms by which astrocytes regulate neural circuits is crucial for elucidating the mechanisms underlying cognitive and behavioral abnormalities in psychiatric disorders and for exploring potential therapeutic or intervention strategies. Viral tracers are powerful vehicles for comprehending the complex relationship between neurons and astrocytes. Despite adeno-associated virus 1 (AAV1) based anterograde trans-neuronal tools have been successfully applied for identifying astrocytes connected with presynaptic neurons, analogous retrograde trans-neuronal tools for labeling astrocytes connected to postsynaptic neurons remain under development. Here, we first demonstrate that GfaABC1D promoter embedding AAV11 can transfer retrogradely from axons of postsynaptic neurons to astrocytes. Furthermore, we established an AAV11-based dual-viral recombination strategy for labeling astrocytes connected to specific postsynaptic neurons. Finally, by employing targeted neuronal ablation technology to ablate neurons in a specific brain region, we observed that injection of AAV11 into the downstream brain region of this targeted area exhibited minimal labeling of astrocytes functionally associated with the ablated neurons. This finding indicates that AAV11 holds utility for deciphering pathological alterations in disease-related astrocyte-neuron connectivity networks. This work fills a key gap in tools used for studying astrocyte-neuron connectivity networks.

## Introduction

Astrocytes, the most abundant glial cells in the mammalian brain, are increasingly recognized as critical regulators of brain development and physiology, primarily through dynamic and bidirectional interactions with neuronal synapses [1-3]. The concept of bidirectional communication between astrocytes and neurons is embodied in the tripartite synapse hypothesis, which emphasizes that astrocytes are key elements in synaptic function [4]. Astrocytes achieve efficient and reliable coordination with neuronal synapses through their proximity to the synaptic cleft, specifically via their peri-synaptic processes, thereby enabling bidirectional communication with both presynaptic and postsynaptic neurons [5, 6].

Their functional abnormalities or incorrect connections with neurons are associated with the development of many diseases, including autism spectrum disorder (ASD) [7, 8], epilepsy [9-11], schizophrenia [12, 13], obsessive-compulsive disorder (OCD) [14], and others. Understanding the complex relationships between astrocytes and neural circuits is crucial for elucidating the mechanisms underlying cognitive and behavioral abnormalities in psychiatric disorders and for exploring potential therapeutic or intervention strategies. However, mapping and manipulating astrocyte-neuron connectivity networks has become an important challenge due to the lack of available techniques.

Viral tracers are powerful vehicles for comprehending neuronal circuits and neuron-astrocyte connection [15-18], among which AAV viruses are widely used in neuroscience and gene therapy eliciting stable, long-term gene expression and minimal pathogenic side effects [19-21]. AAV1 and AAV9 exhibit anterograde trans-synaptic tracing capabilities, enabling their application in mapping input-defined functional neural circuits [22]. AAV1 based anterograde trans-neuronal tools have been successfully applied for identifying astrocytes connected with specific presynaptic neurons [20, 22], however, analogous retrograde trans-neuronal tools for labeling astrocytes connected to specific postsynaptic neurons remain under development. Our prior studies have demonstrated that AAV11 exhibits efficient retrograde targeting of projection neurons and facilitates astrocyte-directed transduction [17]. Consequently, this vector holds potential for astrocyte targeting via retrograde trans-neuronal trafficking.

Here, we first demonstrate that GfaABC1D promoter embedding AAV11 can transfer retrogradely from axons of postsynaptic neurons to astrocytes. Moreover, we established an AAV11-based dual-viral recombination strategy for labeling astrocytes connected to specific postsynaptic neurons. Finally, by employing targeted neuronal ablation technology to ablate neurons in a specific brain region, we observed that injection of AAV11 into the downstream brain region of this targeted area exhibited minimal labeling of astrocytes functionally associated with the ablated neurons. This finding indicates that AAV11 holds utility for deciphering pathological alterations in disease-related astrocyte-neuron connectivity networks. This work fills a key gap in tools used for studying astrocyte-neuron connectivity networks.

## Results

### AAV11 can transfer retrogradely from axonal terminals of postsynaptic neurons to astrocytes

Astrocytes exert critical regulatory roles in neuronal circuit function; however, there remains a paucity of tools or vector platforms enabling direct investigation of the functional interplay between neuronal circuits and astrocytes. Accordingly, we screened natural AAV serotypes to identify vectors capable of mediating direct retrograde transduction of astrocytes via trans-neuronal trafficking. Specifically, we investigated whether the natural AAV11 serotype mediates retrograde transduction of astrocytes within injection site-upstream brain regions. We injected the engineered AAV11-GfaABC1D-EGFP into the caudate-putamen (CPu) region at a dosage of 2 × 10^9^ vector genomes (VG) with a volume of 200 nL (Fig. 1A). Three weeks post-injection, brain tissues were harvested, sectioned and imaged. We found a broad spread of EGFP expression in situ and upstream of CPu (Fig. 1B and C), including the basal lateral amygdala (BLA) and the substantia nigra (SN). Subsequently, we performed immunofluorescence staining using anti-GFAP antibodies on the labeled cells within the SN, demonstrating that these labeled cells were predominantly astrocytes (Fig. 1D).

**Figure 1.**
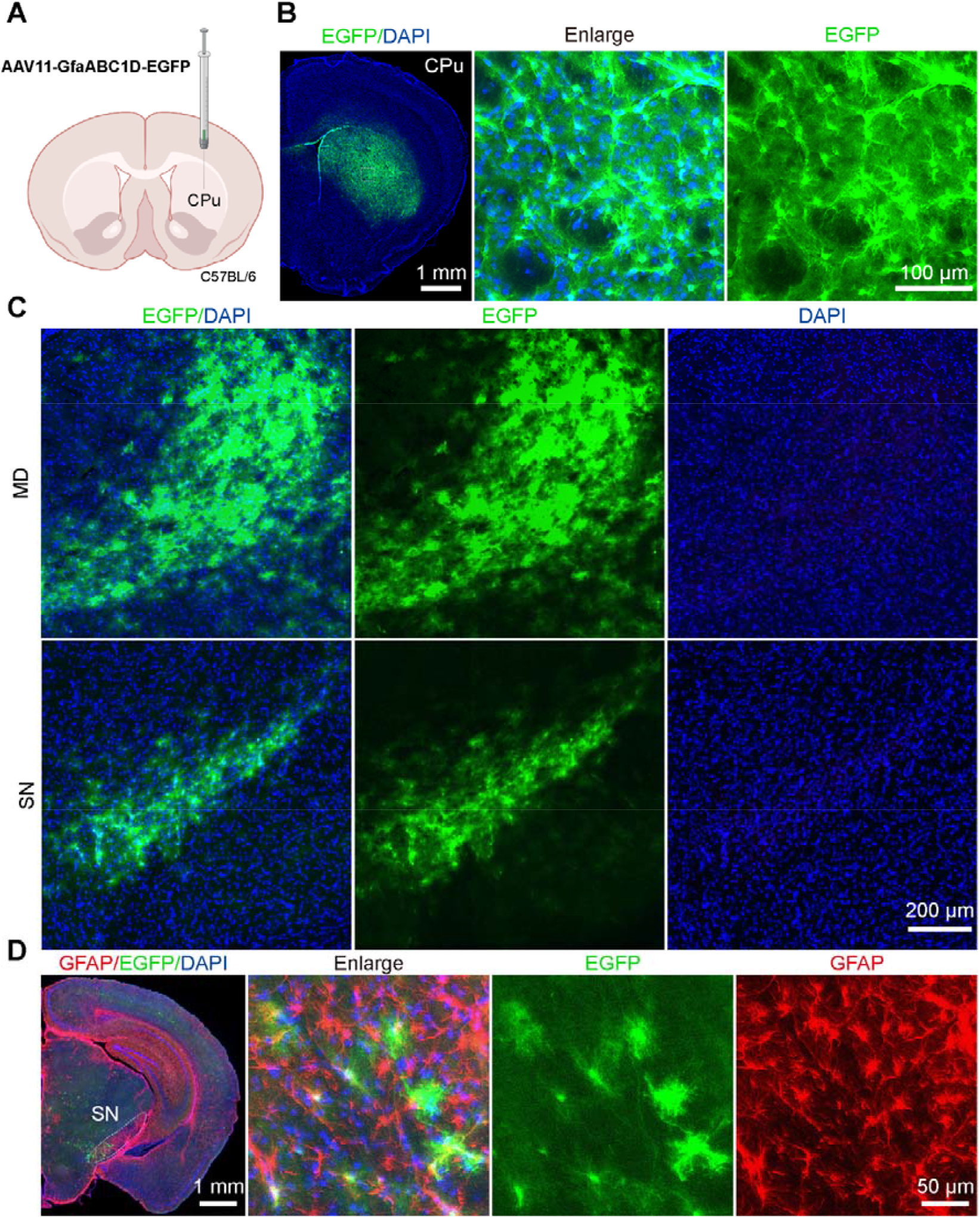
AAV11 can transduce astrocytes distant from the CPu region. A) Schematic diagram of virus injection. AAV11-GfaABC1D-EGFP virus (2 × 10^9^ VG) was injected into CPu area of C57BL/6 mice. B) Representative images showing AAV11 expression in the injection site. Scale bar = 1 mm (left 1 panel), 100 μm (right 2 panels). C) Representative images showing AAV11 expression in the upstream regions of CPu. Scale bar = 200 μm. D) Immune fluorescence imaging of SN region using GFAP antibodies. Scale bar = 1 mm (left 1 panel), 50 μm (right 3 panels). The schematic diagram was created with BioRender.com.

Next, we conducted analogous experiments in other brain regions to validate the broader applicability of this phenomenon. As illustrated in Fig. 2A, we injected the AAV11 vector into the dentate gyrus (DG) of the dorsal hippocampus. Three weeks later, mice were subjected to transcardial perfusion, followed by tissue sectioning and imaging. Consistent with previous reports, abundant AAV-transduced astrocytes were detected in the dorsal hippocampus (co-dHPC) at the injection site (Fig. 2B). Additionally, EGFP-labeled astrocytes were observed in brain regions directly connected to the hippocampus (Fig. 2C), including the medial septal complex (MSC), contralateral hippocampus, and entorhinal cortex (ENT). Subsequently, we performed immunofluorescence staining using anti-GFAP antibodies on the labeled cells within the co-dHPC and ENT, demonstrating that these labeled cells were predominantly astrocytes (Fig. 2D)

**Figure 2.**
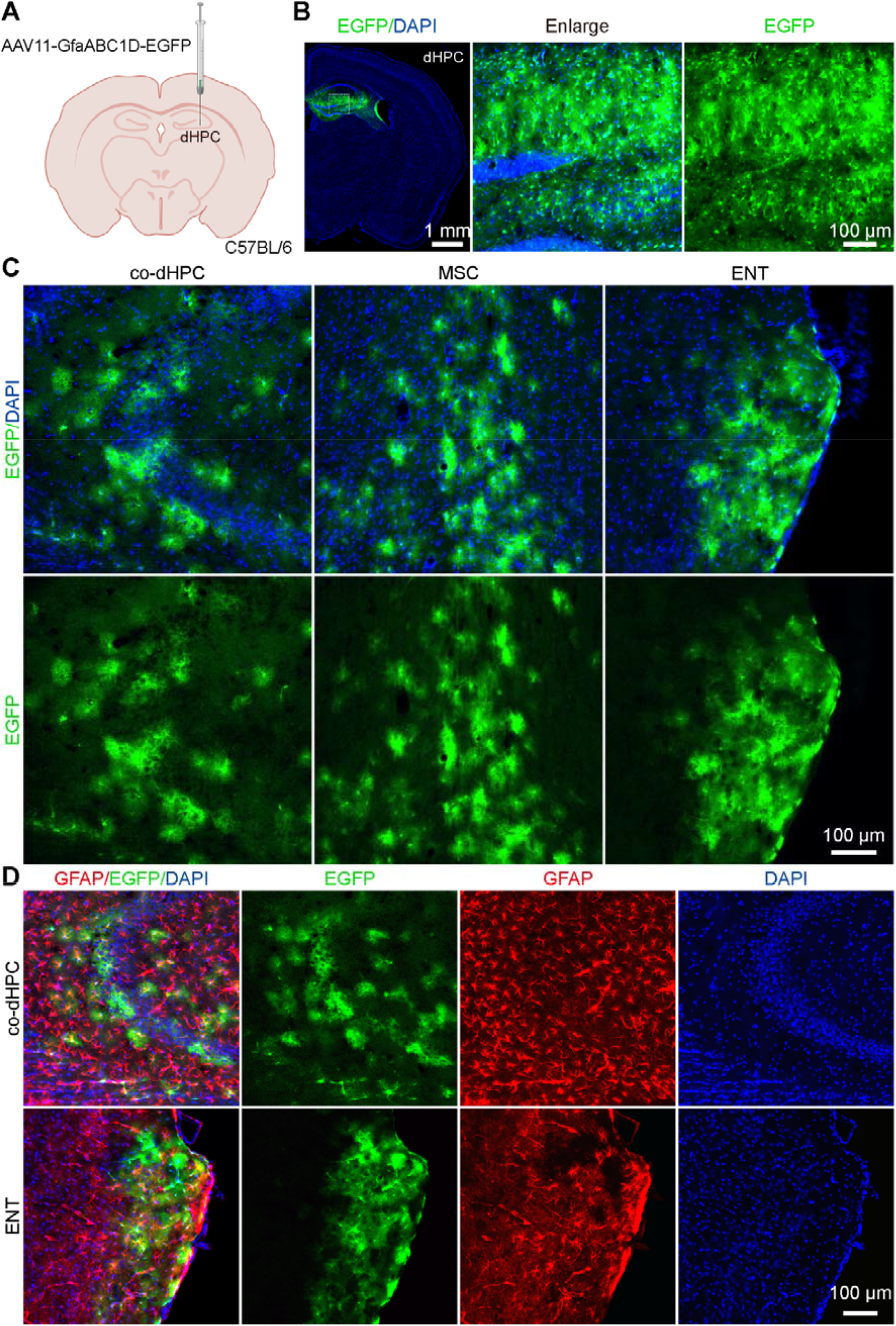
AAV11 can transduce astrocytes distant from the DG region. A) Schematic diagram of virus injection. AAV11-GfaABC1D-EGFP virus (2 × 10^9^ VG) was injected into DG area of C57BL/6 mice. B) Representative images showing AAV11 expression in the DG site. Scale bar = 1 mm (left 1 panel), 100 μm (right 2 panels). C) Representative images showing AAV11 expression in the upstream regions of DG. Scale bar = 200 μm. D) Immune fluorescence imaging of co-dHPC and ENT regions using GFAP antibodies. Scale bar = 100 μm. The schematic diagram was created with BioRender.com.

Certain AAV serotypes, such as AAV1 and AAV9, exhibit anterograde trans-synaptic spread, enabling their widespread propagation [22]. To investigate whether AAV11 mediates anterograde trans-synaptic spread and transduces astrocytes in downstream regions of the injection site, we injected AAV11-GfaABC1D-EGFP into the primary visual cortex (V1) of C57BL/6 mice (Fig. 3A). Three weeks after injection, robust EGFP fluorescence signals were detected at the injection site (Fig. 3B). However, no EGFP-expressing cell bodies were observed in known downstream regions that receive direct projections from V1, including the superior colliculus (SC) and the CPu (Fig. 3C and D), indicating that AAV11 does not mediate anterograde trans-synaptic spread to efficiently transduce astrocytes in downstream regions of the injected site.

**Figure 3.**
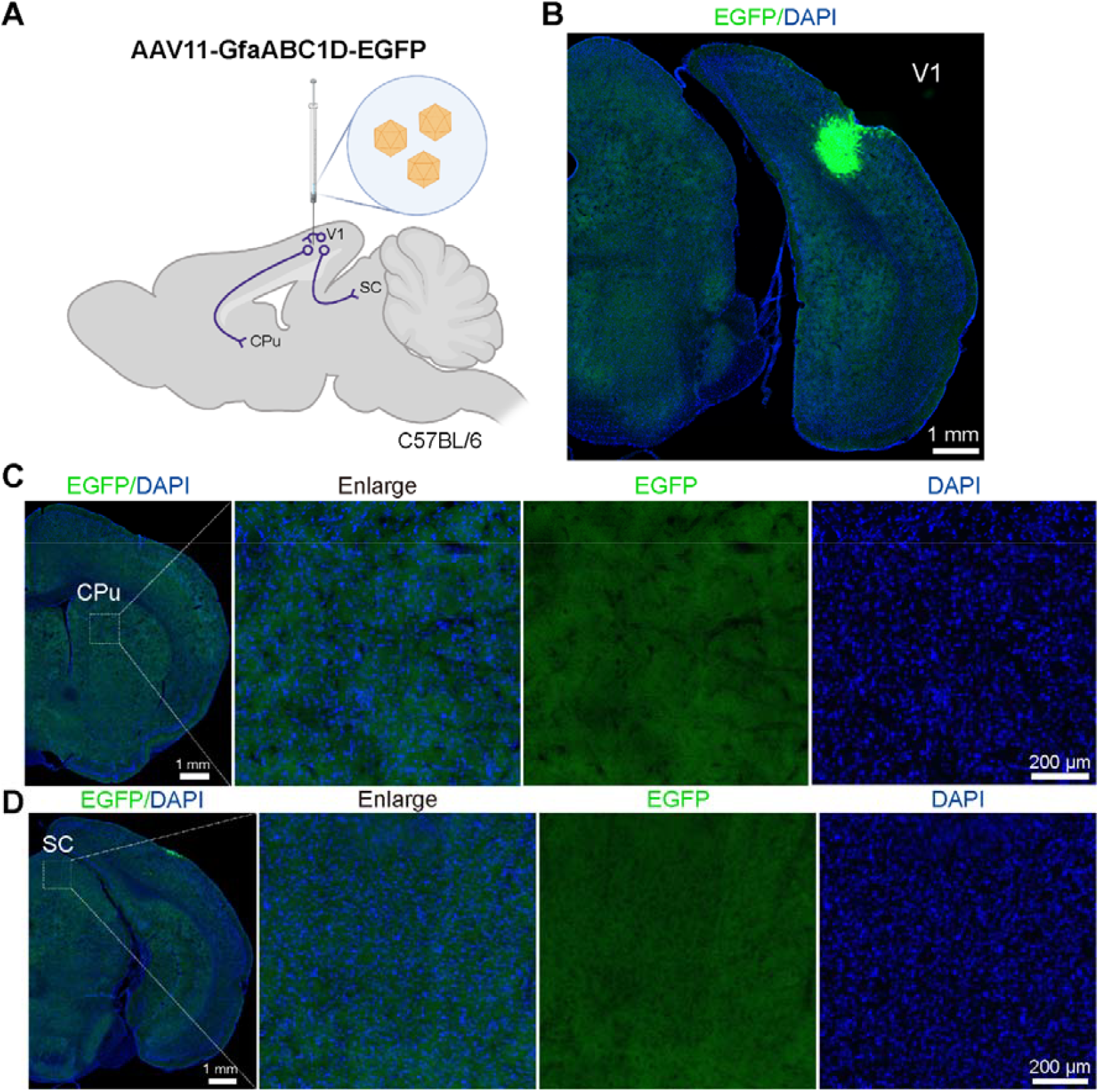
AAV11 does not mediate anterograde trans-synaptic spread to transduce astrocytes. A) Schematic diagram of virus injection. AAV11-GfaABC1D-EGFP virus (2 × 10^9^ VG) was injected into V1 area of C57BL/6 mice. B) Representative images showing AAV11 expression in the V1 site. Scale bar = 1 mm. C) Fluorescence imaging of direct projections from the V1 cortex to CPu brain areas. No EGFP-positive astrocytes were detected in the CPu. Scale bar = 1 mm (left 1 panel), 100 μm (right 3 panels). D) Fluorescence imaging of direct projections from the V1 cortex to SC brain areas. No EGFP-positive astrocytes were detected in the SC. Scale bar = 1 mm (left 1 panel), 100 μm (right 3 panels). The schematic diagram was created with BioRender.com.

These results suggest that AAV11 can transfer retrogradely from axonal terminals of postsynaptic neurons to associated astrocytes.

### An AAV11-based dual-viral recombination strategy for labeling astrocytes connected to specific postsynaptic neurons

To label astrocytes functionally connected to specific postsynaptic neurons, we developed an AAV11-based dual-viral recombination strategy. By combining AAV11 with the ribozyme-activated mRNA trans-ligation system (StitchR) [23], we constructed two vectors—AAV11-GfaABC1D-EGFP-3.0 and AAV11-GfaABC1D-EGFP-4.0—each carrying a fragment of EGFP (Fig. 4A). Fluorescent labeling occurs only when both vectors co-localize within the same cell and express their respective exogenous gene fragments, enabling reconstitution of functional EGFP. For the experimental group, AAV11-GfaABC1D-EGFP-3.0 (2× 10^9^ VG) and AAV11-GfaABC1D-EGFP-4.0 (2× 10^9^ VG) were separately injected into the CPu and SN of the same mouse (Fig. 4B). For the control group, AAV11-GfaABC1D-EGFP-3.0 was injected into the CPu alone, while AAV11-GfaABC1D-EGFP-4.0 was injected into the SN alone (Fig. 4B). Three weeks post-injection, mice were subjected to transcardial perfusion, followed by tissue sectioning and imaging. Notably, no fluorescent labeling was detected in either the CPu or SN of control mice receiving a single vector alone. In contrast, fluorescent labeling of astrocytes was observed in the SN of experimental mice (Fig. 4C). Subsequently, we performed immunofluorescence staining using anti-GFAP antibodies on the labeled cells within the SN, demonstrating that these labeled cells were predominantly astrocytes (Fig. 4D) These results demonstrated that dissection of astrocytes connected to specific postsynaptic neurons can be implemented by combining AAV11 with StitchR system.

**Figure 4.**
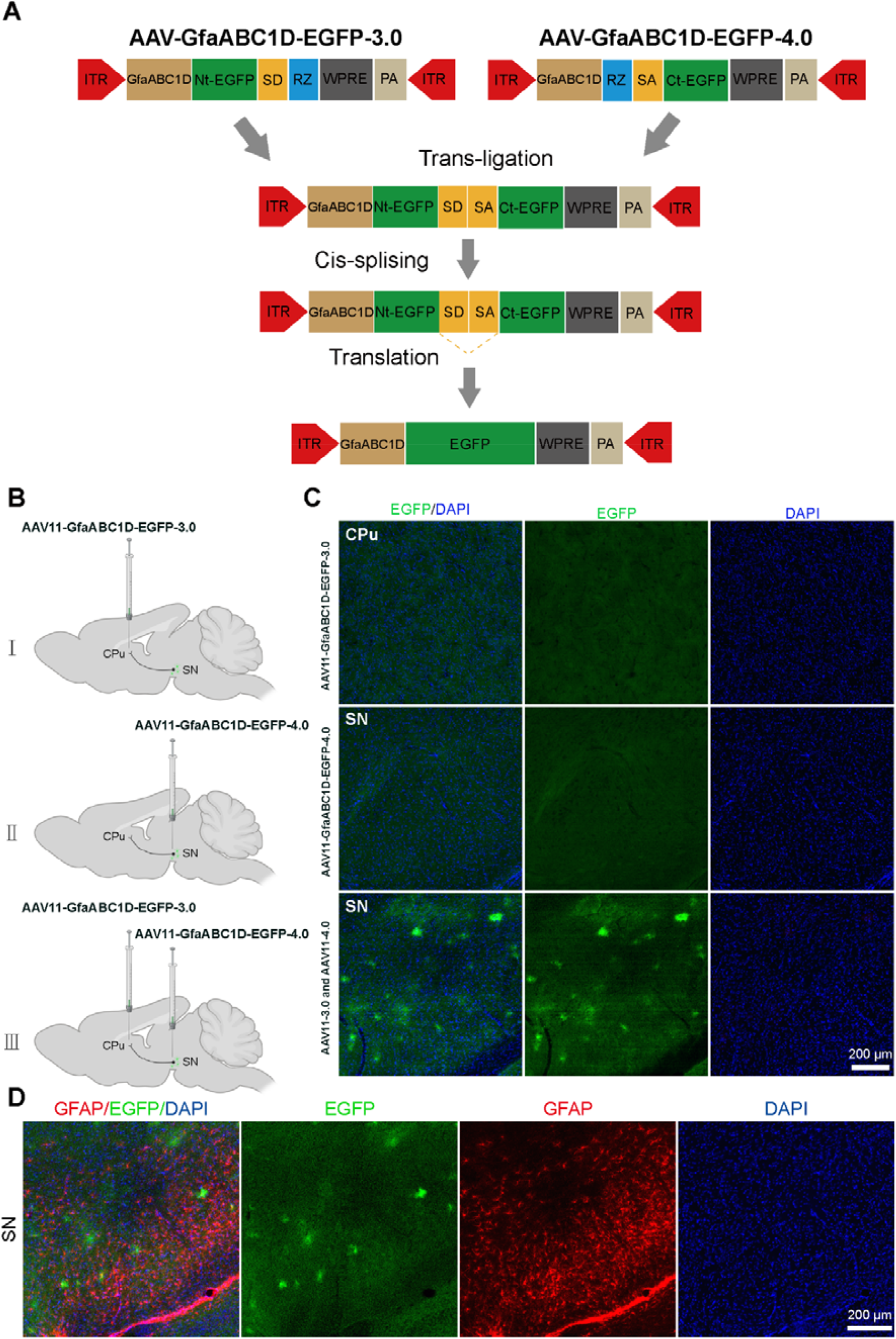
Establishing an AAV11-based dual-viral recombination strategy for labeling astrocytes connected to specific postsynaptic neurons. A) Design of an AAV11_based dual-viral recombination strategy. The ORF of EGFP, containing an intron with splice donor (SD) and splice acceptor (SA) sequences, was split into two non-overlapping N-terminal (Nt) and C-terminal (Ct) vectors. Small catalytic ribozymes (Rz) were utilized at the 3’ end of the Nt-GFP and the 5’ end of the Ct-GFP vectors to create precise RNA termini and achieve scarless trans-ligation. B) Schematic diagram of virus injection. I: AAV11-GfaABC1D-EGFP-3.0 virus (2 × 10^9^ VG) was injected into CPu area of C57BL/6 mice. II: AAV11-GfaABC1D-EGFP-4.0 virus (2 × 10^9^ VG) was injected into SN area of C57BL/6 mice. III: AAV11-GfaABC1D-EGFP-3.0 and AAV11-GfaABC1D-EGFP-4.0 were separately injected into the CPu and SN of the same mouse. C) Representative images showing AAV11 expression. Scale bar = 100 μm. D) Immune fluorescence imaging of SN region using GFAP antibodies. Scale bar = 200 μm. Schematic diagrams were created with BioRender.com.

### AAV11 holds utility for deciphering variation of astrocyte-neuron connectivity networks

To evaluate whether AAV11 can be used for dissecting the variation of astrocyte-neuron connectivity networks, we employed targeted neuronal ablation technology to ablate neurons in a specific brain region, followed by injection of AAV11 into the downstream brain region of this targeted area (Fig. 5A). We injected a mixture of AAV9-EF1a-DIO-DTA (7.5 × 10^8^ VG) and AAV9-hSyn-SV40 NLS-Cre (7.5 × 10^8^ VG) into the SN region (Fig. 5A), and the DTA element expression induced the death of SN neurons. Subsequently, we injected AAV11-GfaABC1D-EGFP (2× 10^9^ VG) into the CPu, the projection areas of the SN region. After three weeks, brain tissues were harvested, sliced and imaged. We observed that injection of AAV11-GfaABC1D-EGFP into the CPu region exhibited minimal labeling of SN astrocytes functionally associated with the ablated neurons (Fig. 5B and C). This finding indicates that AAV11 transduces astrocytes across brain regions through a neuronal circuit-dependent mechanism and holds utility for deciphering pathological alterations in disease-related astrocyte-neuron connectivity networks.

**Figure 5.**
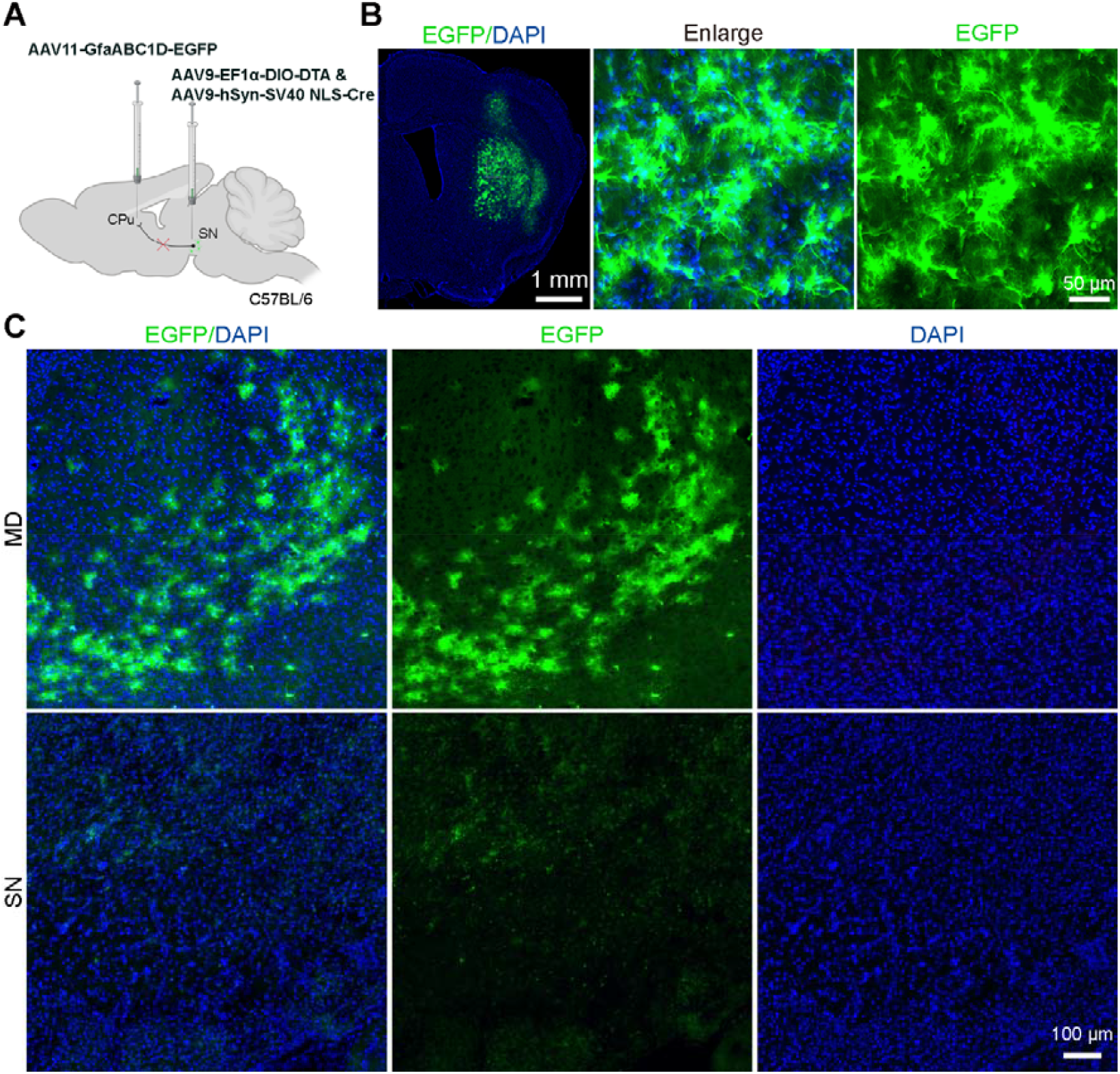
AAV11 can be used to decipher the astrocyte -neuron connectivity networks in disease conditions. A) AAV11-based strategy for astrocyte-neuron labeling in disease models of neuronal ablation. A mixture of AAV9-EF1a-DIO-DTA (7.5 × 10^8^ VG) and AAV9-hSyn-SV40 NLS-Cre (7.5 × 10^8^ VG) were injected to ablate the neurons in the SN. Ten days later, the AAV11-GfaABC1D-EGFP (2× 10^9^ VG) was injected into the CPu. B) Representative images showing AAV11 expression in the injection site of CPu. Scale bar = 1 mm (left 1 panel), 50 μm (right 2 panels). C) Representative images showing AAV11 expression in the upstream regions of CPu. The astrocytes in the midbrain dopaminergic (MD) region, which are upstream projections of the CPu brain area, show normal GFP labeling signals, while the astrocytes in the substancia nigra (SN) region, which are also upstream projections of the CPu brain area, do not show normal GFP labeling signals. Scale bar = 100 μm. The schematic diagram was created with BioRender.com.

## Discussion

AAV is widely recognized as one of the safest and most versatile viral vectors for DNA delivery, enabling efficient gene overexpression, knockdown, and editing [24]. It has been extensively applied to neural circuit tracing and mapping [25], in vivo fluorescence imaging [26], disease model establishment [27, 28], and evaluation of molecularly targeted therapeutic strategies for neurological disorders [29]. For targeted delivery to the central nervous system (CNS), common AAV delivery routes include local intraparenchymal injection [30], intravenous administration [21, 31], anterograde trans-synaptic trafficking [21], and retrograde transport via absorption at axon terminals [32]. Herein, we demonstrate that AAV11 enables retrograde trans-neuronal targeting of astrocytes via projection neurons—thus expanding its utility in studying astrocyte-neuron connectivity networks.

Astrocytes, the most abundant glial cell type in the brain, are crucial for maintaining the homeostasis and health of the brain [33, 34]. Despite the traditional description of astrocytes as supportive partners for neurons, they are now recognized as active participants in the development and plasticity of dendritic spines and synapses [35]. Notably, recent studies have highlighted that astrocyte dysfunction is a common feature across multiple neurodegenerative diseases such as Alzheimer’s disease and Parkinson’s disease [36-38]. Consequently, astrocytes have emerged as key therapeutic targets for these disorders. For instance, in vivo astrocyte-to-neuron conversion represents a promising neuroregeneration strategy for neurodegenerative disease intervention [39]. Specifically, in vivo viral delivery of short hairpin RNA targeting PTB (shPTB) or RNA-targeted CasRx has been shown to efficiently convert astrocytes into functional neurons, thereby alleviating symptoms of neurodegenerative diseases [40, 41]. Our findings show that AAV11 enables retrograde transduction of astrocytes across neurons via trans-neuronal trafficking—this vector thus offers a novel tool for developing therapeutic interventions targeting diseases driven by dysregulated neuron-astrocyte interactions.

Most AAVs requires the GPR108 and AAVR (KIAA0319L) receptors for cellular entry, with the exceptions of AAV4 and rh32.33, which depend on GPR108 but are independent of AAVR [42-44]. Given that AAV11 and rh32.33 both belong to the AAV4 lineage clade [45], it is possible that AAV11 depends on GPR108 for successful cellular transduction. Notably, recent studies have shown that AAV11 additionally depends on the AAVR2 receptor for this process [46]. The efficient retrograde trans-neuronal transduction of astrocytes mediated by AAV11 may be associated with GPR108 or AAVR2; however, this remains a hypothesis that requires validation through additional experiments.

### Experimental Section

#### Plasmids construction and AAV vector manufacturing

To construct the transfer plasmid pAAV-GfaABC1D-EGFP-WPRE-pA, GfaABC1D sequence was synthesized and inserted into the pAAV-hSyn-EGFP-WPRE-pA digested with MluI and NcoI restriction endonucleases (Thermo Fisher Scientific, Waltham, MA, USA). AAV packaging plasmids were obtained following the construction method of pAAV2/11 [17]. AAV viruses were packaged with the helper plasmid pAd-DeltaF6 (Addgene, Watertown, MA, USA, 112867), pAAV2/11 [17] and pAAV-GfaABC1D-EGFP-WPRE-pA.

HEK-293T cells (American Type Culture Collection, Manassas, VA, USA) were cultured in suspension using Balance CD medium (Chuangling Cell-wise, Shanghai, China, CW01001) supplemented with 1% penicillin/streptomycin (BasalMedia, Shanghai, China, S110JV), and kept at 37 □ in a 5% CO2 atmosphere. Sixteen hours before the transfection process, these cells were shifted to DMEM (BasalMedia, L110KJ) containing 2% fetal bovine serum (Thermo Fisher Scientific GIBCO, Waltham, MA, USA, 10099141C) and 1% penicillin/streptomycin (BasalMedia, S110JV). Briefly, AAV vectors were generated through a transient transfection method utilizing three plasmids and linear polyethylenimine (Polysciences, Warrington, PA, USA, 24765-1) for the process. At 72 hours following transfection, the viral particles were collected, followed by purification using an iodixanol gradient ultracentrifugation technique, and then concentrated and exchanged into PBS containing 0.001% Pluronic F68 (Thermo Fisher Scientific, Waltham, MA, USA, 24040032) via Amicon® Ultra centrifugal filters (Merck Millipore, Billerica, MA, USA, UFC910024). The titers of the purified recombinant AAVs were determined by quantitative PCR using the iQ SYBR Green Supermix kit (Bio-Rad, Hercules, CA, USA, 1708884) with primer WPRE-F (5’-ATGCCTTTGTATCATGCTATTGCT-3’) and WPRE-R (5’-CACGGAATTGTCAGTGCCCAA-3’). These viral vectors were then divided into aliquots and stored at -80 □ for future use.

### Research animals

All procedures conducted in this study were approved (approval No. APM20026A) by the Animal Care and Use Committee of the Innovation Academy for Precision Measurement Science and Technology, Chinese Academy of Sciences. Adult male C57BL/6 mice (provided by Hunan SJA Laboratory Animal Company, Changsha, Hunan, China) aged 8-10 weeks were used for the experiments. The mice were housed in a controlled environment with a 12/12-hour light/dark cycle within specific pathogen-free facilities. The temperature was maintained between 22°C and 24°C, and the humidity levels were kept between 40% and 60%. Both water and food were provided *ad libitum* to the mice.

### Stereotaxic AAV injection

Mice were deeply anaesthetized using 1% pentobarbital intraperitoneally (i.p., 50 mg/kg body weight) and placed in a stereotaxic apparatus (Item: 68025 - stereotaxic apparatus and 68030 - mice adaptor, RWD, China). The injection coordinates were selected according to Paxinos and Franklin’s *The Mouse Brain in Stereotaxic Coordinates*, 4th edition [47]. A small volume of virus was injected into the CPu (relative to bregma: anterior-posterior-axis (AP) +0.80 mm, medial-lateral-axis (ML) ±2.00 mm, and dorsal-ventral-axis (DV) –3.30 mm), DG (relative to bregma: AP –2.15 mm, ML ±1.30 mm, and DV –2.00 mm), and SN (relative to bregma: AP –3.1 mm, ML ±1.20 mm, and DV –4.30 mm) and V1 (relative to bregma: AP –3.90□mm, ML□±□2.6□mm, and DV –1.30□mm). Viruses were injected at a rate of 0.03 μL/min using a stereotaxic injector equipped with a pulled glass capillary (Stoelting, Wood Dale, IL, USA, 53311). After the injection was complete, the micropipette was held for an additional 10 minutes before being withdrawn. Animals were allowed to recover from anaesthesia on a heating pad. Three weeks after injection, the mice were sacrificed and brain tissues were collected via transcardic infusion of PBS and 4% paraformaldehyde solution.

### Immunofluorescence and imaging

Brain slices preparation and imaging were performed according to a previously reported method [48]. The brains were immersed in a 4% paraformaldehyde solution overnight, then dehydrated in a 30% sucrose solution. Subsequently, they were sectioned into 40 μm-thick slices using a microtome (Thermo Fisher Scientific, Waltham, MA, USA), collected in antifreeze solution, and stored at -20 °C until further use. For IBA1 staining, sections were incubated with a primary antibody goat anti-GFAP (1:800, Abcam, Cambridge, MA, USA, ab53554), followed by a secondary antibody donkey anti-goat IgG conjugated with Cy3 (1:400, The Jackson Laboratory, Bar Harbor, ME, USA, 305-165-003). After PBS washing, all brain slices mounted on microscope slides were counterstained with DAPI (1:4000, Beyotime, Shanghai, China) and sealed with 70% glycerol. Imaging was conducted using either the Olympus VS120 Slide Scanner microscope (Olympus, Tokyo, Japan) or Leica TCS SP8 confocal microscope (Leica, Wetzlar, Germany).

## Data Availability Statement

The authors declare that all supporting data for this study are included within the manuscript or can be obtained from the authors.

## Author Contributions

N.L., K.L. and Y.G. contributed to the study idea and design; K.L. and F.X. contributed to funding acquisition and resources; N.L., H.Z., and K.L. performed the experiments and data acquisition; N.L., Y.G., H.Z., and K.L. accomplished data analysis; N.L., K.L., Y.G., H.Z. and F.X. drafted the manuscript, and contributed to review and editing. All authors read and approved the final manuscript.

